# Controlled and Selective Photo-oxidation of Amyloid-β Fibrils by Oligomeric *p*-Phenylene Ethynylenes

**DOI:** 10.1101/2021.11.03.467121

**Authors:** Adeline M. Fanni, Daniel Okoye, Florencia A. Monge, Julia Hammond, Fahimeh Maghsoodi, Tye D. Martin, Gabriella Brinkley, M. Lisa Phipps, Deborah G. Evans, Jennifer S. Martinez, David G. Whitten, Eva Y. Chi

## Abstract

Photodynamic therapy (PDT) has been explored as a therapeutic strategy to clear toxic amyloid aggregates involved in neurodegenerative disorders such as Alzheimer’s disease. A major limitation of PDT is off-target oxidation, which can be lethal for the surrounding cells. We have shown that a novel class of oligo-*p*-phenylene ethynylene-based compounds (OPEs) exhibit selective binding and fluorescence turn-on in the presence of pre-fibrillar and fibrillar aggregates of disease-relevant proteins such as amyloid-β (Aβ) and α-synuclein. Concomitant with fluorescence turn-on, OPE also photosensitizes singlet oxygen under illumination through the generation of a triplet state, pointing to the potential application of OPEs as photosensitizers in PDT. Herein, we investigated the photosensitizing activity of an anionic OPE for the photo-oxidation of toxic Aβ aggregates and compared its efficacy to the well-known but non-selective photosensitizer methylene blue (MB). Our results show that while MB photo-oxidized both monomeric and fibrillar conformers of Aβ40, OPE oxidized only Aβ40 fibrils, targeting two histidine residues on the fibril surface and a methionine residue located in the fibril core. Oxidized fibrils were shorter and more dispersed, but retained the characteristic β-sheet rich fibrillar structure and the ability to seed further fibril growth. Importantly, the oxidized fibrils displayed low toxicity. We have thus discovered a class of novel theranostics for the simultaneous detection and oxidization of amyloid aggregates. Importantly, the selectivity of OPE’s photosensitizing activity overcomes the limitation of off-target oxidation of currently available photosensitizers, and represents a significant advancement of PDT as a viable strategy to treat neurodegenerative disorders.

## Introduction

Developing photosensitizers that are selective for their intended targets would significantly advance their application in photodynamic therapy (PDT) in the treatment of numerous human diseases as off-target oxidation is a major drawback of current PDT technology.^1,2^ Herein, we evaluate the selective photosensitizing activity of a novel class of conjugated polyelectrolytes, oligo-*p*-phenylene ethynylenes (OPEs), recently developed for detecting the aggregated conformations of amyloid proteins. Demonstration of OPE’s selective photo-oxidation of amyloid protein aggregates will enable the development of these compounds in theranostic applications for the simultaneous detection and treatment of protein misfolding diseases, including Alzheimer’s and Parkinson’s diseases.

A major pathological hallmark of Alzheimer’s disease is the deposition of amyloid plaques composed of the amyloid-β (Aβ) peptide,^3,4^ which results from the abnormal misfolding and aggregation of the peptides into small oligomers that subsequently grow into large fibrils.^5-8^ The oligomers, which are transient and heterogeneous in nature, are known to be more neurotoxic than the mature fibrils. The mechanism of their toxicity is still unclear but their interactions with cell membranes leading to membrane destabilization and pore formation have been proposed to cause cell apoptosis.^9-11^ Aβ fibrils also play a key role in neurodegeneration through impairment of axonal transport^12,13^ or by inducing the aggregation of tau protein and seeding the formation of neurofibrillary tangles.^14^ Additionally, amyloid aggregates are also involved in the rapid and predictable spatiotemporal disease progression through cell-to-cell transmission.^15-17^ Because of the central roles amyloid aggregates play in Alzheimer’s disease pathogenesis, their selective degradation and clearance is an attractive therapeutic approach.

Therapeutic strategies targeting amyloid aggregates, specifically Aβ plaques, are currently being investigated and include the use of enzymes (neprilysin^18^, insulin-degrading enzyme^19,20^ and endothelin-converting enzyme^21,22^) and anti-Aβ immunotherapy.^23^ These approaches have several limitations including the non-selective degradation of the native Aβ peptide which is believed to have important physiological functions.^24,25^

PDT is a therapeutic strategy currently used in oncology^26,27^ and dermatology.^28^ In PDT, a photosensitizer is exposed to light, generating singlet oxygen ^1^O_2_ (photosensitization type II) and/or free radicals (photosensitization type I) through energy transfer from an excited triplet state.^29^ Those species then oxidize biological molecules, including proteins, lipids and amino acids, ultimately leading to cell death (e.g., oncology).^30^ A number of photosensitizers have been investigated for the photo-oxidation of Aβ including riboflavin,^31^ rose bengal,^32^ porphyrin-based molecules,^33,34^ flavin-based compounds,^31,35^ polymer^36^ and carbon nanodots,^37,38^ metal complexes^39-41^, fullerene-based materials^42,43^ and methylene blue (MB).^44^ These are non-selective and induce photo-oxidation of both Aβ monomers and aggregates, as well as other biomolecules in the vicinity to the photosensitizers. For example, MB, a major FDA-approved photosensitizer used in oncology^45,46^, has been investigated for photo-oxidation of Aβ plaques both *in vitro* and *in vivo*.^44^ Aside from oxidizing both Aβ monomers and fibrils, MB also non-selectively binds to and oxidize negatively charged proteins, lipids and nucleic acids^46^, leading to cellular apoptosis. These studies show that photo-oxidation of Aβ monomers inhibits fibrillation^31-33^, which could be beneficial but likely also cause the loss-of-function of the peptide. Encouragingly, photo-oxidation of fibrils has been found to disassemble the fibrils into shorter structures displaying lower cellular toxicity,^35^ pointing to the potential that PDT targeted at the aggregated, pathogenic conformation of amyloid proteins can be a beneficial therapeutic approach. Several fibril-specific photosensitizers have recently been developed by Kanai et al., including those based on fibril-binding dyes thioflavin-T^47,48^ and curcumin^49^. Importantly, these studies found that photooxidation caused Aβ aggregate degradation, which attenuated aggregate toxicity and reduced aggregate levels in the brains of AD mouse models.

We recently showed that a class of novel oligomeric *p*-phenylene ethynylene compounds (OPEs) selectively bind to and detect β-sheet rich amyloid fibrils.^50-53^ The small and negatively charged OPE_1_^2-^ (Table 1), characterized by one repeat unit with carboxyethyl ester end groups and sulfonate terminated side chains, detects fibrils made of two model amyloid proteins, insulin and lysozyme,^50,51^ and two disease-relevant proteins, Aβ and α-synuclein.^52^ More importantly, OPE_1_^2-^ is also capable of selectively detecting the more toxic, pre-fibrillar aggregates of Aβ42 and α-synuclein.^52^ OPE’s superior sensor performance compared to that of the commonly used thioflavin T dye could be attributed to its high sensitivity to fluorescence quenching wherein the conjugated sensor is quenched in an aqueous solvent and binding to amyloid aggregates reverses quenching and leads to fluorescence recovery or turn-on.^53-56^ The ability of OPE_1_^2^ to detect a wider set of protein aggregate conformations stems from the combination of different modes that leads to fluorescence turn-on of the OPEs, including hydrophobic unquenching, backbone planarization, and OPE complexation upon binding to amyloid aggregates.^50,51,55^

**Table 1:**
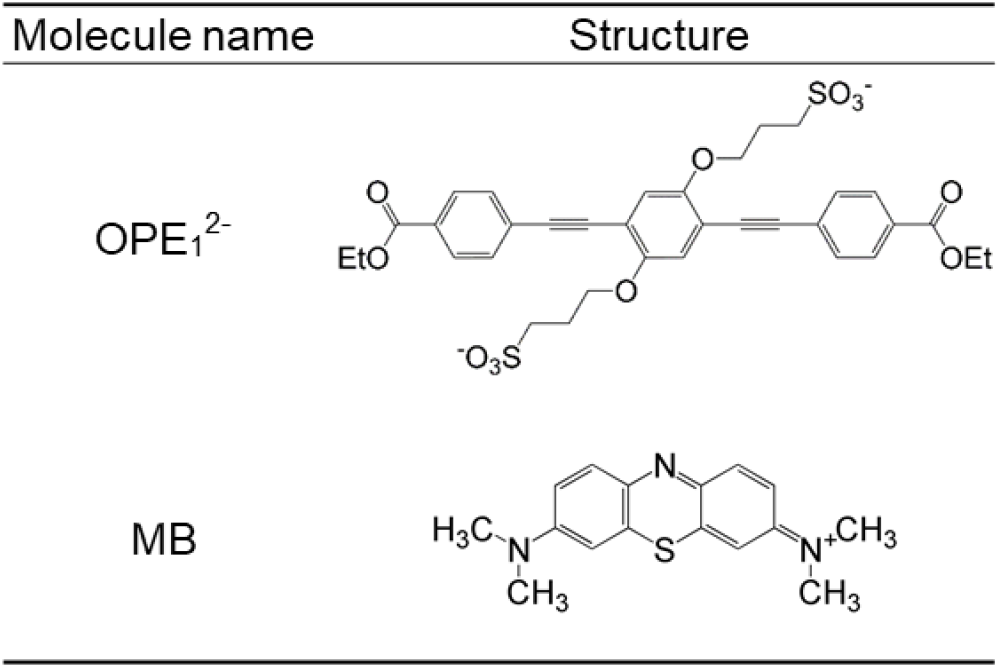
Structures of photosensitizers OPE_1_^2-^ and methylene blue (MB) used in this study

In addition to sensing, we recently showed that fluorescence turn-on of OPE_1_ ^2-^ from binding to the cationic detergent cetyltrimethylammonium bromide (CTAB) is accompanied by the generation of a triplet state which subsequently photosensitizes ^1^O_2_.^57^ Importantly, as the unbound (quenched) state of OPE_1_ ^2-^ does not have photosensitizing activity, both fluorescence and photosensitizing properties of the OPE are selective, which makes this probe highly promising as a photosensitizer that is simultaneously controllable with light and selective for the pathogenic conformations of amyloid proteins. As photo-oxidation can potentially reduce the toxicity and promote the clearance of pathogenic amyloid aggregates, the simultaneous sensing and oxidation of the aggregates by OPEs may represent a novel theranostic approach to simultaneously detect and treat protein misfolding diseases.

In this study, we evaluated the potential of OPE_1_^2-^ as a selective photo-oxidizer for Aβ fibrils over its monomeric counterpart and compared its activity to the well-known but non-specific photosensitizer MB. Oxidation of both Aβ monomers and fibrils with light exposure in the presence of OPE_1_^2-^ or MB was characterized. Oxidized amino acids on Aβ fibrils were identified and quantified, and the effect of fibril oxidation on fibril morphology, secondary structures, cell toxicity, and fibril seeding potency were evaluated.

## Results

### Spectroscopic features of OPE_1_^2-^ and MB in the presence of Aβ40 monomers and fibrils

We have previously shown that the photosensitization activity of OPE_1_^2-^ is activated by its fluorescence turn-on through complexation with oppositely charged detergent molecules.^57^ In this study, we first measured the absorbance and fluorescence spectra of both OPE_1_^2-^ and MB in the presence of monomeric or fibrillar Aβ40. Aβ40 fibrils were produced by incubating the monomeric peptide at 150 µM in pH 8.0 40 mM Tris buffer at 37 °C for 23 days. At this pH, the Aβ40 peptide has a net charge of -4.4.^58^ The unincubated Aβ does not show any features on TEM images (Figure 1A) indicating that the peptide is most likely monomeric, while incubated peptide showed large clusters of fibrils (Figure 1B). These fibrils were previously characterized to be rich in β-sheets while the monomeric peptides are mainly random coils.^52^

**Figure 1:**
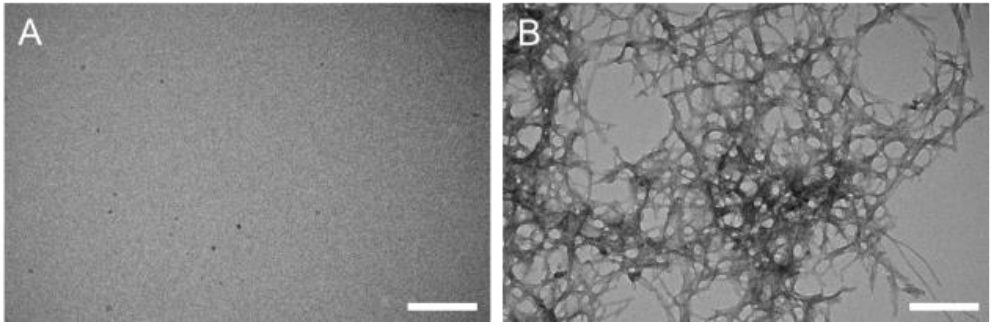
TEM images of Aβ monomers (A) and fibrils (B). Freshly solubilized Aβ monomers do not show any features on TEM images while after 23 days of incubation, Aβ formed large clusters of fibrils. Scale bar = 200 nm.

OPE_1_^2-^ and MB absorbance and emission spectra were measured in the presence of different concentrations of Aβ40 monomers and fibrils (Figure 2A1-2A2 and 2B1-2B2). OPE^12^-absorbance spectrum is characterized by two main peaks at 316 and 368 nm. In the presence of Aβ monomers or fibrils, OPE^12^-absorbance spectra were unchanged in terms of peak positions and intensities, consistent with previous results.^52^ The absorption spectrum of MB, characterized by one main peak at 663 nm, increased by 14-22% in the presence of either Aβ monomers or fibrils, but the peak shape and position stayed unchanged.

**Figure 2:**
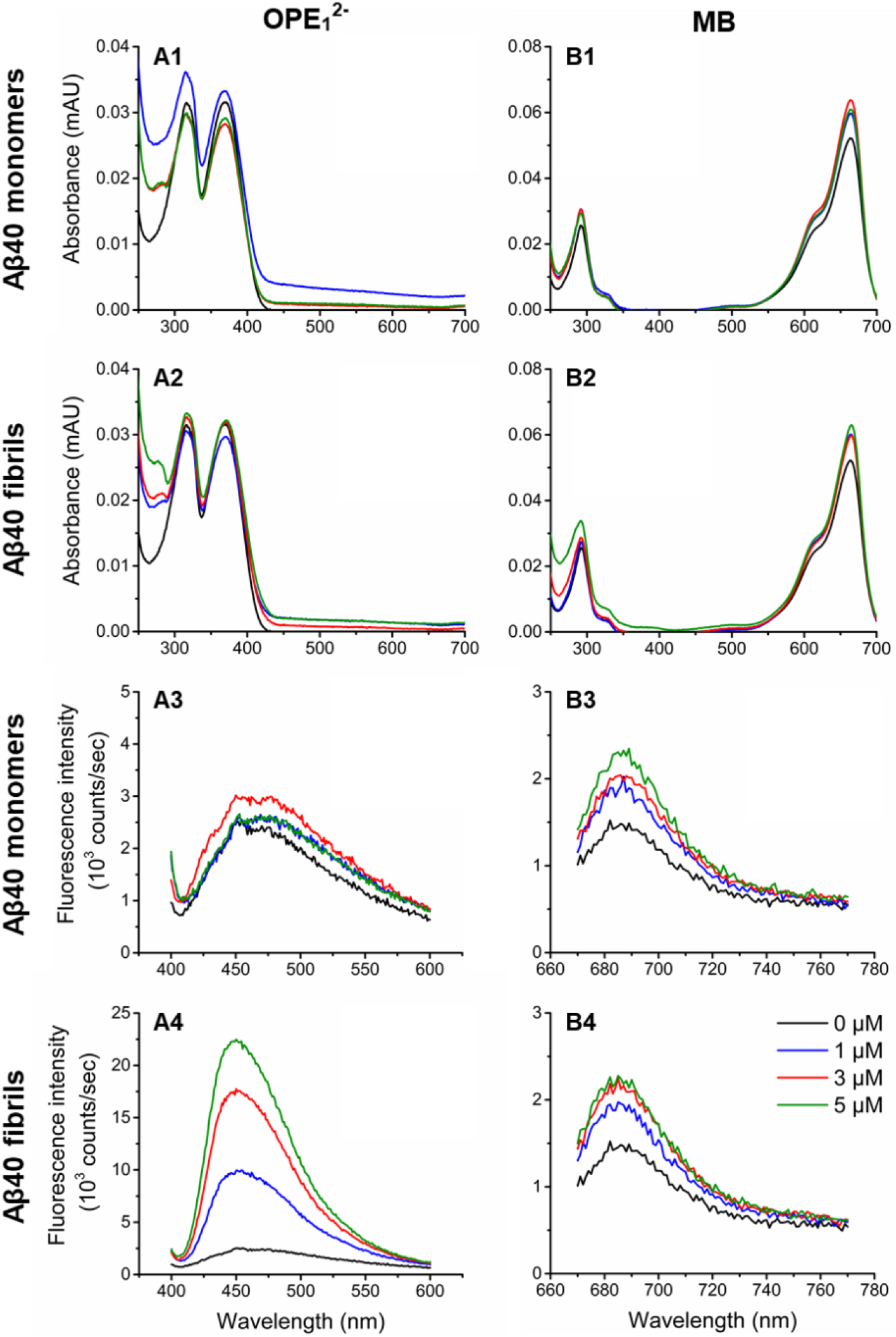
OPE_1_^2-^ displays a selective fluorescence turn-on in the presence of Aβ40 fibrils, while MB exhibits weak fluorescence increases in the presence of both monomeric and fibrillar Aβ40. Absorbance (1 and 2) and fluorescence emission spectra (3 and 4) of OPE_1_^2-^ (A) and MB (B) at 1 µM in the presence of varying concentrations of Aβ40 monomers or fibrils (0, 1, 3 and 5 µM are shown in black, blue, red and green, respectively).

OPE_1_^2-^ and MB fluorescence were also characterized (Figure 2A3-2A4 and 2B3-2B4). At 1 µM, OPE_1_^2-^ is quenched in water and in the presence of 1 to 5 µM Aβ40 monomers, no significant change in emission intensity was observed (Figure 2A3). However, when added to 5 µM fibrillar Aβ40, OPE_1_^2-^ fluorescence emission drastically increased more than 5-fold blue shifted (Figure 2A4). These results indicate that OPE_1_^2-^ displays a selective fluorescence turn-on upon binding to Aβ40 fibrils. MB is also a fluorescent molecule characterized by an emission peak centered at 685 nm. In the presence of both monomeric and fibrillar Aβ40, MB emission moderately increased by 30-50% (Figure 2B3-2B4), which could be due to its interaction to the negatively charge Aβ peptides causing MB to planarize, which prevents non-radiative relaxation decay of the photo-excited MB.^44^ The similar levels of fluorescence increases indicate that MB interacts similarly to both monomeric and fibrillar Aβ conformations, making its interaction nonselective.

### DNPH dot blot of Aβ40 oxidation

Having confirmed that OPE_1_^2-^ selectively binds amyloid fibrils over monomeric Aβ peptides, we then determined if OPE_1_^2-^ also only photo-oxidizes Aβ40 fibrils and not the monomers. The oxidation states of both Aβ40 conformers irradiated in the presence of OPE_1_^2-^ were characterized and compared to MB by monitoring the carbonyl content of the peptide using the DNPH (2,4-dinitrophenylhydrazine) dot blot assay.^59^ The assay was carried out on 5 µM Aβ40 fibrils and monomers incubated for up to 6 hours either in the dark or with irradiation in the presence or absence of 1 µM photosensitizer (Figure 3). In the dark (Figure 3A), no photo-oxidation is detected in Aβ monomers incubated alone (column 1) or with a photosensitizer (columns 4 and 7) as no dots are visible. Aβ40 fibrils showed faint dots (Figure 3A, column 2) that did not become darker over time, indicating a low level of oxidation might have occurred during the 23 days of incubation to produce fibrils. The low intensity dots could also be due to non-specific binding of DNPH to the dense fibrils. Light exposure alone also did not cause any oxidation of Aβ40 monomers or fibrils as no changes in their carbonyl contents were observed (Figure 3B, columns 1 and 2). However, in the presence of both photosensitizers and under light irradiation, Aβ40 fibrils quickly became oxidized, as evidenced by large increases in carbonyl content where the dots clearly became darker with irradiation time (Figure 3B, columns 5 and 8). Importantly, Aβ monomers irradiated in the presence of OPE did not become photo-oxidized as no dots were visible (Figure 3B, column 4). Monomers irradiated in the presence of MB showed small increases in carbonyl content as faint dots were visible (Figure 3B, column 7). Taken together, the dot blot assay shows that MB oxidized both monomers and fibrils, although to different extents, and OPE_1_^2-^ only oxidized Aβ fibrils under irradiation.

**Figure 3:**
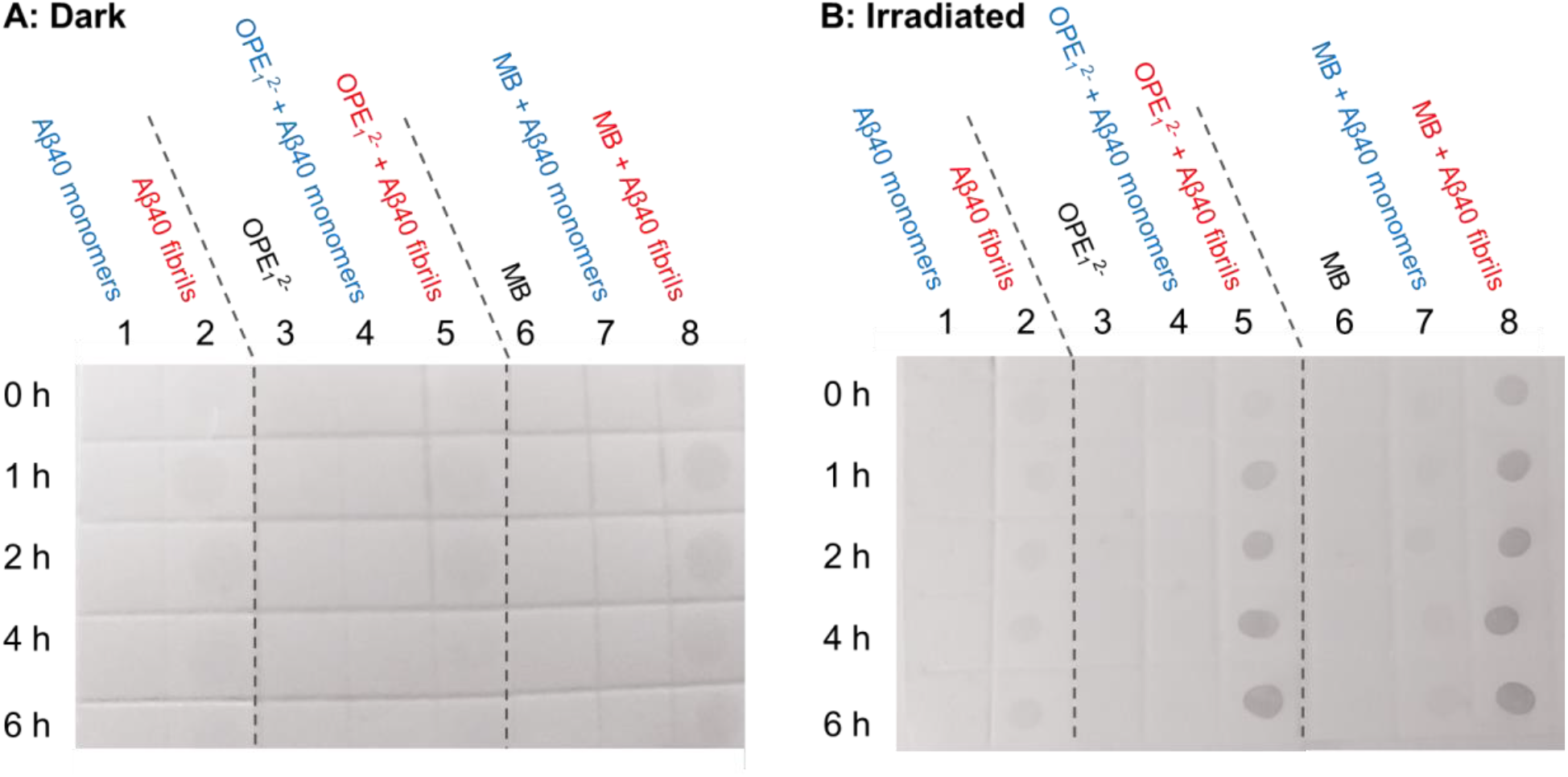
Results from DNPH dot blot assay show that OPE_1_^2-^ selectively oxidized Aβ40 fibrils over monomers under light irradiation while MB non-selectively oxidized both oxidized Aβ40 fibrils and monomers. Aβ monomers (5 µM) and fibrils (5 µM) in the presence of OPE_1_^2-^ (1 µM) or MB (1 µM) at different incubation times in the dark (A) or under light irradiation (B). Dots indicate carbonyl groups of oxidized amino acids and the darker the dots, the higher the carbonyl content. Note that at 0 h, Aβ fibrils incubated with MB (column B8) showed a dot, which could be due to short light exposure during membrane preparation causing MB to photosensitize the oxidation of the fibrils.

### Mass spectrometry characterization of Aβ40 oxidation

The DNPH dot blot assay qualitatively showed that OPE_1_^2-^ selectively induced photo-oxidation of Aβ fibrils, but not monomers. To confirm this selectivity, we turned to molecular techniques of electrospray ionization mass spectrometry (ESI-MS). Figure 4 shows ESI-MS spectra of Aβ40 monomer (25 µM) before and after irradiation in the presence of OPE_1_^2-^ or MB (1 µM). A quadruple charged ion (m/z of 1082) was observed for the Aβ40 monomer without irradiation (Figure 4A), which corresponds to monomer’s expected and deconvolved mass of 4329. The ESI-MS analysis of Aβ40 monomer obtained after irradiation in the presence of OPE_1_^2-^ gave results similar to those obtained from the native Aβ40 monomer (Figure 4B), indicating that irradiation in the presence of OPE_1_^2-^ did not induce any mass changes in the peptide. In contrast, m/z peaks of Aβ monomers irradiated in the presence of MB (Figure 4C) showed a mixture of oxidized species and their corresponding mass increases (oxidation of the following amino acids are suspected: His (155 g/mol) into dehydro-2-imidiazolone derivative (169 g/mol), Met (149 g/mol) into sulfoxide (165 g/mol), and Tyr (181 g/mol) into 3,4-dihydroxyphenalalanine (197 g/mol))^49,60,61^). Oxidation of Aβ monomers was additionally confirmed by reverse phase HPLC (Figure S1) where the oxidized peptide eluted earlier in a broad peak revealing the presence of a mixture of more hydrophilic peptides; this result is consistent with the presence of oxidized peptides that are more hydrophilic. Our results are also consistent with previous reports on MB induced oxidation of monomeric Aβ.^44^

**Figure 4:**
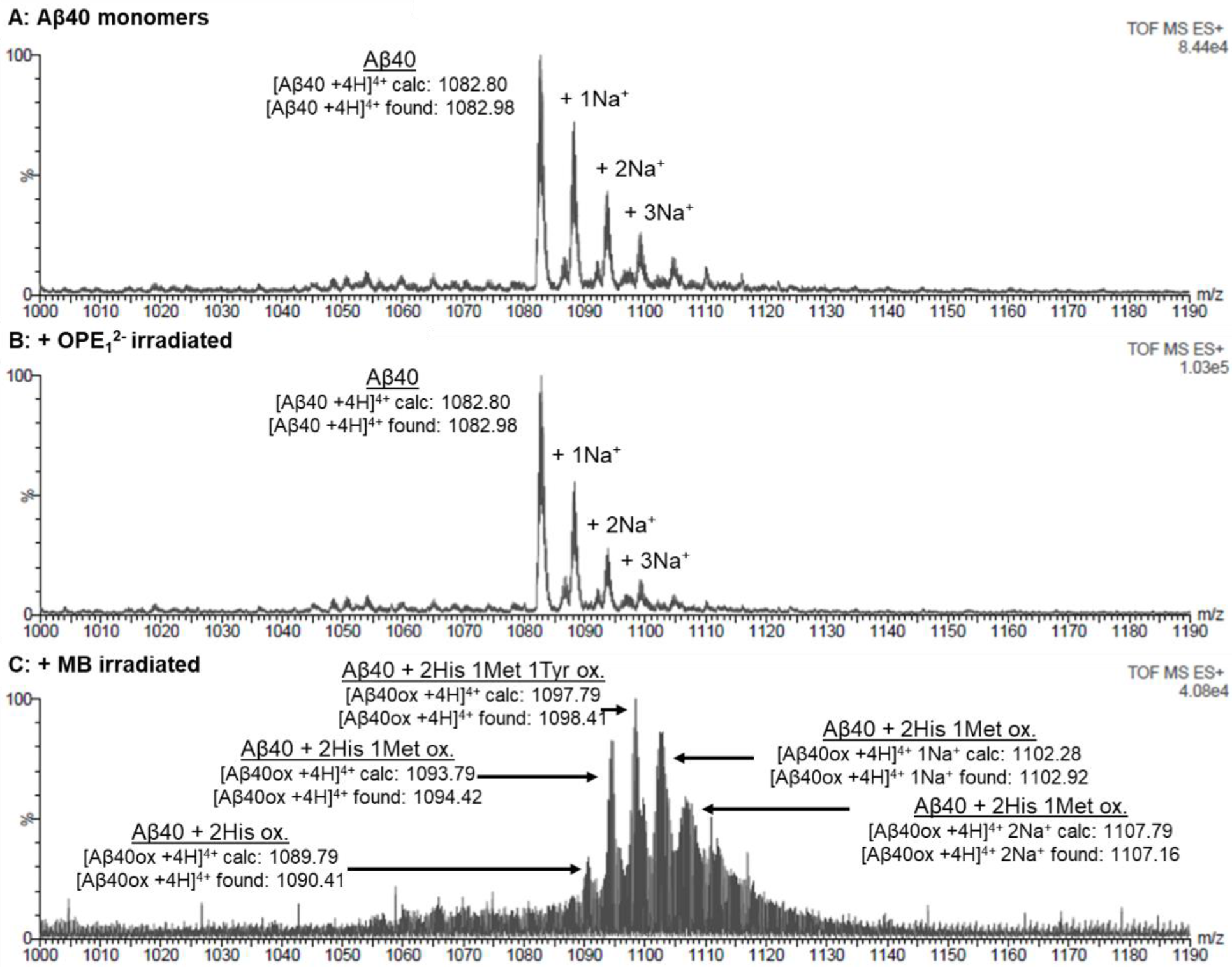
Mass spectrometry confirms that while MB induces photo-oxidation of Aβ monomers, OPE_1_^2-^ does not. Electrospray ionization mass spectrometry (ESI-MS) chromatograms of 25 µM of Aβ40 monomers (A) and Aβ40 monomers irradiated for 4 hours in the presence of 1 µM of OPE_1_^2-^ (B) or MB (C). Non-irradiated Aβ40 monomer is characterized by a m/z peak at 1082.80 corresponding to [AB40 +4H]^4+^. A similar profile was found after irradiation in the presence of OPE_1_^2-^ indicating that OPE did not oxidize the monomeric peptide. After irradiation of Aβ40 in the presence of MB, a mixture of oxidized monomers with higher m/z values appeared.

The oxidation of Aβ40 fibrils by both photosensitizers was also characterized by ESI-MS. Fibrils were first solubilized by enzymatic digestion with Endoproteinase LysC (Figure 5A) and the fragments were separated by HPLC and analyzed with mass spectrometry; out of the three fragments, only the Aβ29-40 fragment which contains the oxidizable Met35 residue was well-resolved by mass spectrometry. The expected mass of the native and oxidized Aβ29-40 fragments are 1084.6 Da and 1100.6Da, respectively (Figure 5B). The size of these fragments in the presence of the Na adduct is shown in Figure 5B. After irradiating 25 µM Aβ40 fibrils in the presence of 1 µM MB, all Aβ29-40 peaks shifted by 16 Da (Figure 5D) indicating complete oxidation of the fragment. Met35 is likely oxidized as it is the only photo-oxidizable amino acid in this fragment^35^. After irradiating the fibrils with OPE_1_^2-^, oxidized peaks were observed but the non-oxidized fragments were still present in the sample (Figure 5E). OPE_1_^2-^ thus only partially oxidized Met35 located in the core of Aβ fibrils while MB completely oxidized the residue.

**Figure 5:**
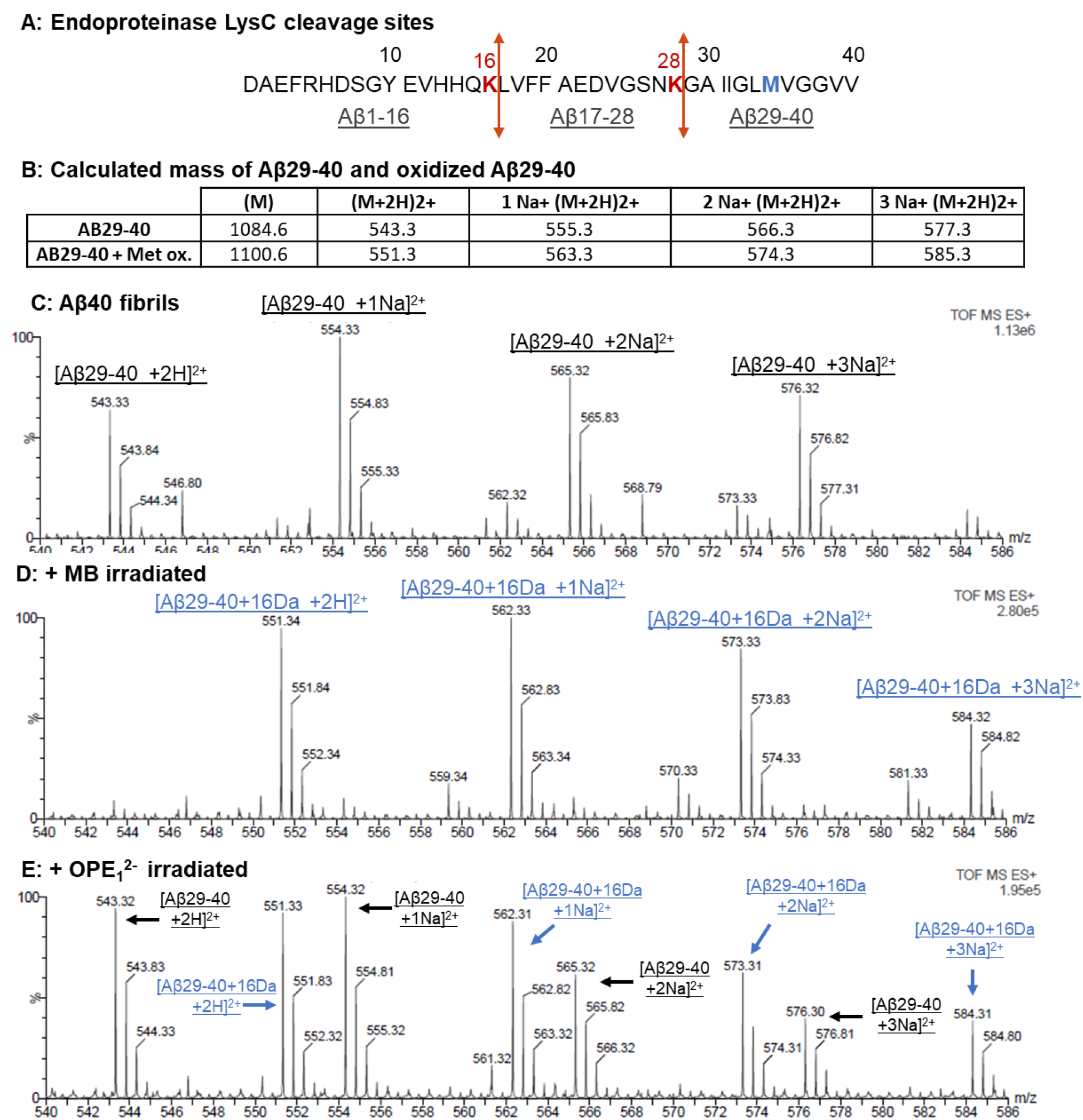
Mass spectrometry confirms that both OPE_1_^2-^ and MB induces photo-oxidation of Aβ fibrils. A: Cleavage sites of Endoproteinase LysC in Aβ40 peptide as indicated in red. Cleavage product Aβ29-40 is the only well-resolved fragment from mass spectrometry. In blue is the suspected oxidized methionine in Aβ29-40 fragment. B: Calculated masses of the Aβ29-40 fragment with methionine and oxidized methionine. C-E: ESI mass spectrometry chromatograms of the Aβ29-40 fragment generated from non-irradiated fibrils (C) and after 4 hours of irradiation in the presence of 1 µM MB (D) or OPE_1_^2-^ (E).

### Amino acid analysis of oxidized Aβ40

Amino acid analysis (AAA) was carried out to further confirm Aβ40 oxidation as well as to more completely identify oxidized amino acids (Figure 6). Aβ40 fibrils before and after light irradiation in the presence of either OPE_1_^2-^ or MB were analyzed. As we have confirmed that OPE_1_^2-^ does not oxidize Aβ monomers (Figures 3 and 4), only Aβ monomers irradiated in the presence of MB was analyzed (Figure 6A). In AAA, Aβ peptides were cleaved into individual amino acids by hydrolysis using 6 N HCl. Note that Met can become oxidized during acid hydrolysis^62^. For this reason, no conclusion was made regarding the effect of MB and OPE_1_^2-^ on methionine content from this analysis.

**Figure 6:**
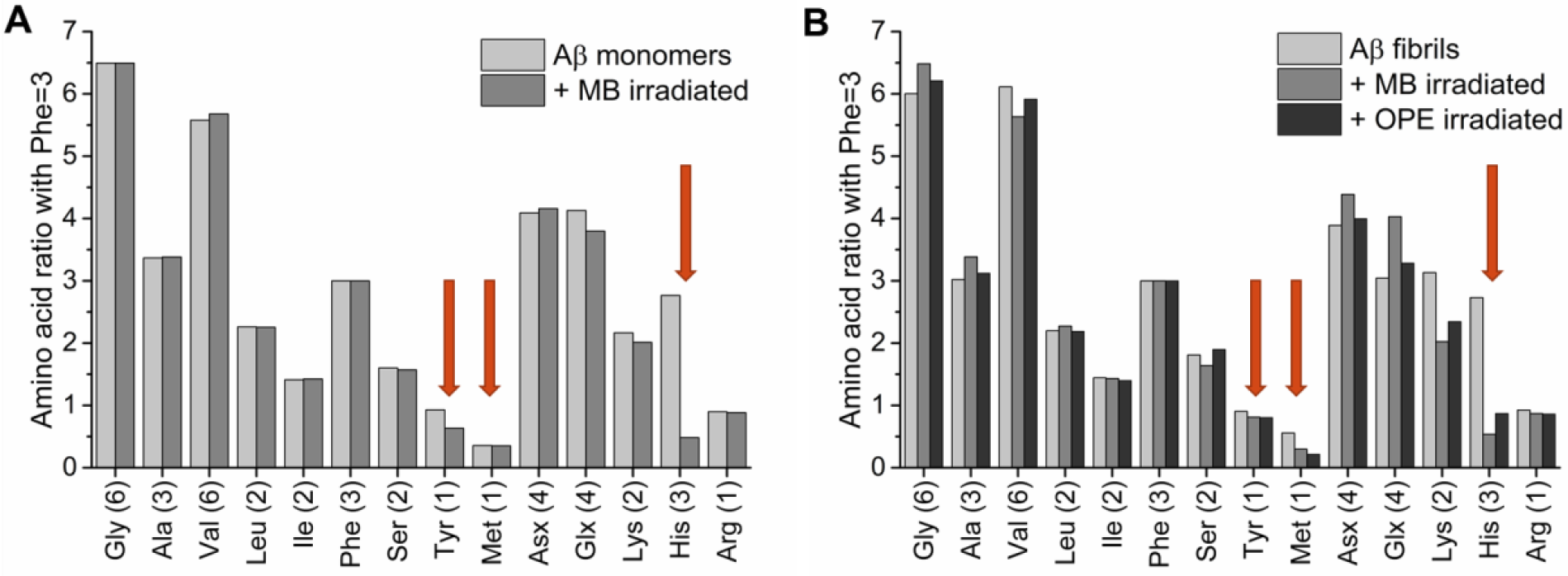
His, Tyr, and Met residues are oxidized in Aβ monomers by MB, while His and Met residues are oxidized in Aβ fibrils by both MB and OPE_1_^2-^. Amino acid analysis of 25 µM Aβ40 monomers (A) and fibrils (B) before and after 4-hour irradiation in the presence of 5 µM of MB or OPE_1_^2-^. Amino acid content was determined by normalizing the raw data with known signal generated for phenylalanine. The x axis describes the type and number of amino acids expected in a single Aβ40 peptide. Glutamic acid and glutamine could not be differentiated by this analysis and they together appear as Glx. Similarly, aspartic acid and asparagine appear as Asx. Ile content is lower than expected (around 1.2 instead of 2), which might be explained by poor hydrolysis of the Ile-Ile bond. Red arrows indicate the three amino acids that can be photo-oxidized – Tyr, Met and His. Note that as Met can be partially oxidized during hydrolysis, the effect of MB and OPE_1_^2-^ on Met content cannot be conclusively made with this method.

AAA results show that the His content in Aβ40 monomers and fibrils irradiated in the presence of MB reduced from 3 to 1 (Figure 6A and 6B). Thus 2 out of the 3 His residues were oxidized by MB. Interestingly, Tyr content in Aβ40 monomers was partially reduced, which is supported by the ESI-MS analysis (Figure 4), but stayed unchanged in Aβ40 fibrils after irradiation with MB. Aβ40 fibrils irradiated in the presence of OPE_1_^2-^ or MB (Figure 6B) showed reduced His content and no change in Tyr content. Combined with ESI-MS results, it can thus be concluded that two His, one Tyr, and one Met residues were oxidized in Aβ40 monomers but only two His and one Met residues were oxidized in Aβ40 fibrils. The difference is that Tyr is oxidized only in monomers, but not in Aβ fibrils, which could be due to differences in the residue’s solvent exposure in the two different Aβ conformations. Tyr oxidation has been found to depend on solvent exposure^63^. For example, Tyr residues in insulin were found to be non-oxidized in the native hexameric conformation but oxidized in an 8 M urea denatured state^64^. Denaturation increases solvent accessibility and thereby the activity of a photosensitizer. In the disordered Aβ monomer, Tyr is more accessible to MB photosensitization whereas in the fibrillar state, Aβ fibril has a core that is inaccessible to the solvent, reducing photo-oxidation of Tyr residues by both MB and OPE.

### Molecular dynamics (MD) simulations of OPE_1_^2-^ binding sites on Aβ40 protofibrils

Having confirmed that OPE_1_2-selectively oxidized 2 out of 3 His residues and partially oxidized the Met residue on Aβ fibrils, we then sought to understand the photo-oxidation pattern of this photosensitizer by analyzing binding sites of OPE on the fibril and their proximity to oxidizable residues. It is known that singlet oxygen species generated by a photosensitizer can only diffuse through 100-200 Å in a biological system^27,65,66^ and their reactivity additionally depends on their environment such as solvent accessibility.^67^ We utilized all-atom MD simulations to identify OPE binding sites on an Aβ protofibril and analyzed the distances of the oxidizable residues to the bound OPEs.

Aβ40 contains 5 amino acids that can be photo-oxidized: His6, His13, His14, Tyr10 and Met35 (Figure 7A). A protofibril made of 24 Aβ9-40 peptides was obtained from the 2LMN^68^ Protein Data Bank (PDB) file (Figure 7C). The first 8 amino acids remained disordered in the fibril and were not represented in this structure. The structure of a disordered Aβ9-40 monomer was obtained by first removing a peptide from the protofibril PDB structure and then performing energy minimization and equilibration for 100 ns (Figure 7B). The locations of oxidizable amino acids within the monomer and protofibril are shown in Figure 7B and 7C, respectively. All 4 oxidizable amino acids of Aβ9-40 are freely exposed to the solvent when the peptide is monomeric. In the protofibril, Met35 is buried inside the β-sheet core while His13, His14 and Tyr10 are located on the surface of the protofibril. His6 is in the disordered N-terminal region and not represented on these structures of Aβ9-40.

**Figure 7:**
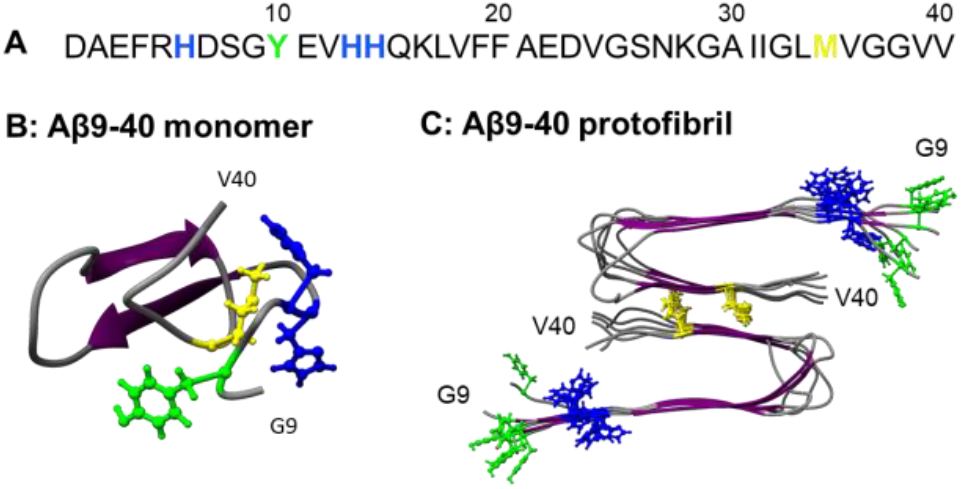
Oxidizable amino acids in the Aβ40 sequence (A) and their locations in an Aβ9-40 monomer (B) and protofibril (C). Met, Tyr, and His residues are shown in yellow, green, and blue, respectively.

OPE_1_^2-^ binding sites on the Aβ40 protofibril were analyzed by MD with 12 OPE molecules placed around the protofibril. After 100 ns of simulation, 11 out of 12 OPE became bound to the protofibril at six different binding sites either as single OPE molecules (sites 1, 4 and 5) or as OPE complexes (sites 2, 3 and 6) (Figure 8A). Three of the binding sites were located on the β-sheet rich protofibril surface (sites 2, 3 and 6; Figure 8B and C), one was located on the β-turn (site 5; Figure 8D), and the last two were at the ends of the protofibril (sites 1 and 4; Figure 8B and E). Tyr and His residues were located within 4 Å of OPE_1_^2-^ in binding sites 2, 3, 5 and 6, which make them highly susceptible for oxidation by singlet oxygen. In contrast, Met was buried inside the core of the fibril which makes it further away from singlet oxygen generated by surface bound OPEs and possibly less prone to oxidation due to its reduced solvent accessibility.^67^ The closest bound OPE_1_^2-^ at site 4 on one end of the protofibril was 5.6 Å away from Met35. The other bound OPEs were further away from Met (between 19.8 – 41.6 Å), which might explain the partial oxidation of Met35 observed.

**Figure 8:**
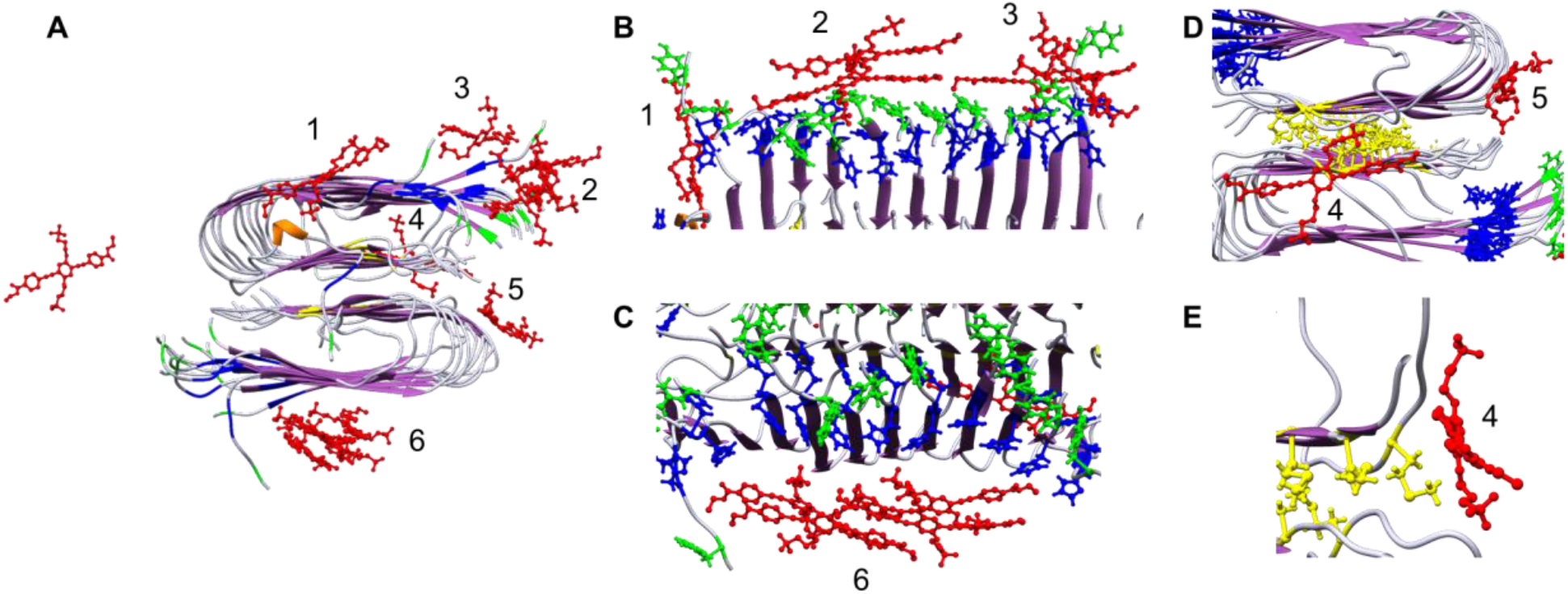
Positions of 12 OPEs (red) placed around an Aβ protofibril after 100 ns of all atom MD simulation (A) show 11 OPEs bound to the protofibril at 6 binding sites. B, C, D, E and F show different zoomed-in views of the 6 binding sites where OPEs bind both as single OPEs or complexes of OPE. Methionine (M), tyrosine (Y) and histidine (H) are shown in yellow, green and blue, respectively. The remainder of the protofibril is shown in ribbon representation with β-sheets colored purple and random coil in white.

### Effect of oxidation on Aβ40 fibril morphology and secondary structures

For PDT to be a viable treatment option, the structural and functional properties of the photo-oxidation products need to be characterized to ensure that they will not be harmful. The effect of oxidation on the fibril’s morphology and secondary structures were assessed by TEM imaging and circular dichroism (CD) spectroscopy, respectively. As shown in Figure 9A, untreated Aβ40 fibrils were long and in large clusters. After irradiation in the absence of a photosensitizer, the fibril clusters appeared smaller, but were still present (Figure 9B). When the fibrils were irradiated with either MB or OPE_1_^2-^, the clusters were largely gone and shorter fibrils were observed (Figure 9C and D).

**Figure 9:**
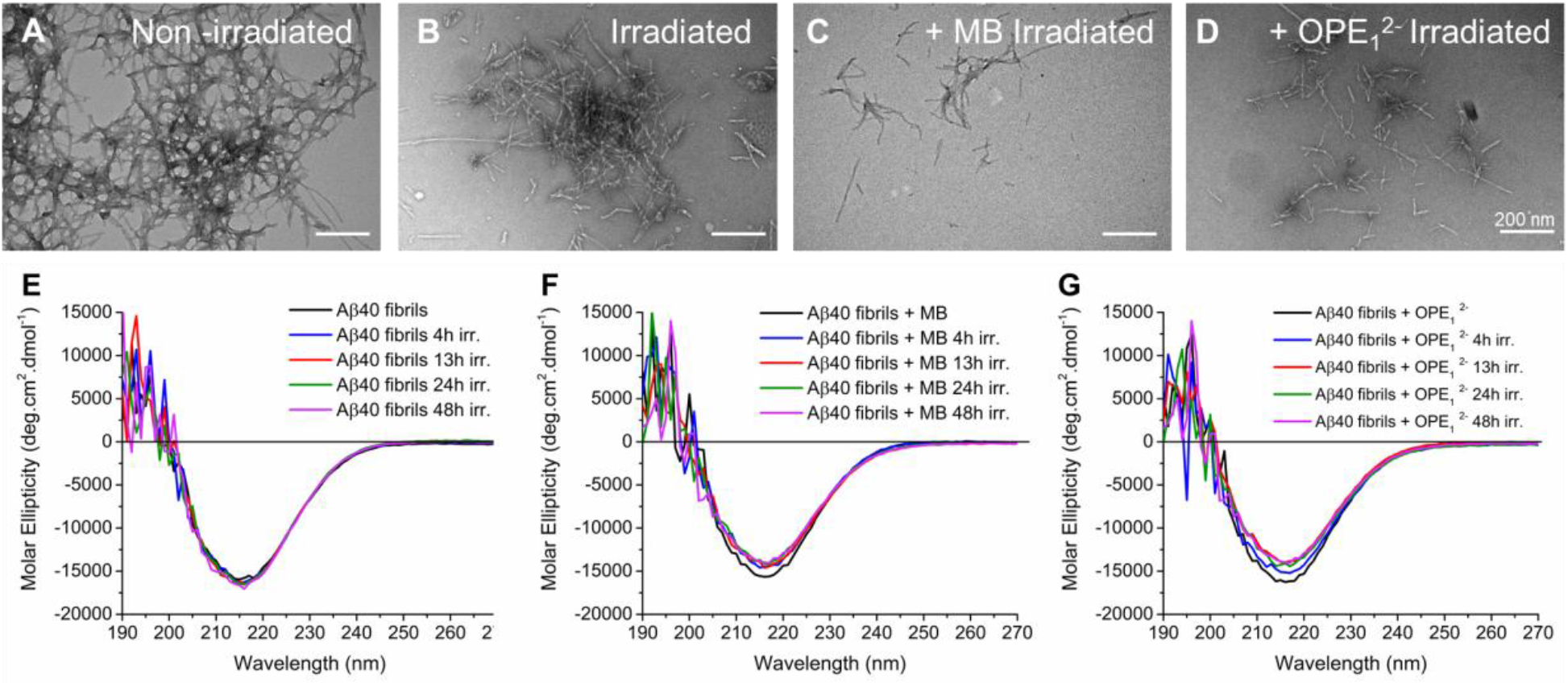
Photo-oxidation led to some breakdown of fibrils, but did not alter their secondary structures. TEM images and CD spectra of Aβ40 fibrils irradiated in the absence or presence of a photosensitizer. TEM images of 5 µM Aβ40 fibrils: non-irradiated (A), irradiated for 4 hours (B), irradiated in the presence of MB for 4 hours (C), and irradiated in the presence of OPE_1_^2-^ for 4 hours (D). CD spectra of 50 µM Aβ fibrils irradiated for various times: irradiated alone (E), irradiated with 10 µM MB (F), and incubated 10 µM OPE_1_^2-^ (G).

CD spectra of Aβ40 fibrils showed positive and negative peaks at 192 nm and 215 nm, respectively, indicating a β-sheet rich structure. Irradiation of the fibrils did not affect their secondary structures (Figure 9E). Irradiation in the presence of OPE_1_^2-^ or MB (Figure 9G and H) also did not result in significant changes to the fibril’s secondary structures; the shorter fibrils remained rich in β-sheets. Taken together, our results indicate that photo-oxidation induced some breakdown of fibrils, but did not alter their β-sheet rich secondary structures.

### Effect of oxidation on Aβ40 fibril seeding potency

The ability to seed aggregation is a prominent property of amyloid fibrils as fibril elongation by the addition of monomers is likely the primary aggregation pathway for Aβ40.^69-71^ The effect of oxidation on the seeding potency of Aβ fibrils was evaluated by monitoring Aβ40 monomer (50 µM) aggregation in the presence of 2.6 µM of non-oxidized and photo-oxidized fibril seeds after 72 hours of incubation at 37°C (Figure 10). Fibril seeds were prepared by sonicating mature Aβ40 fibrils in a bath sonicator for 10 minutes (Figure S2).

**Figure 10:**
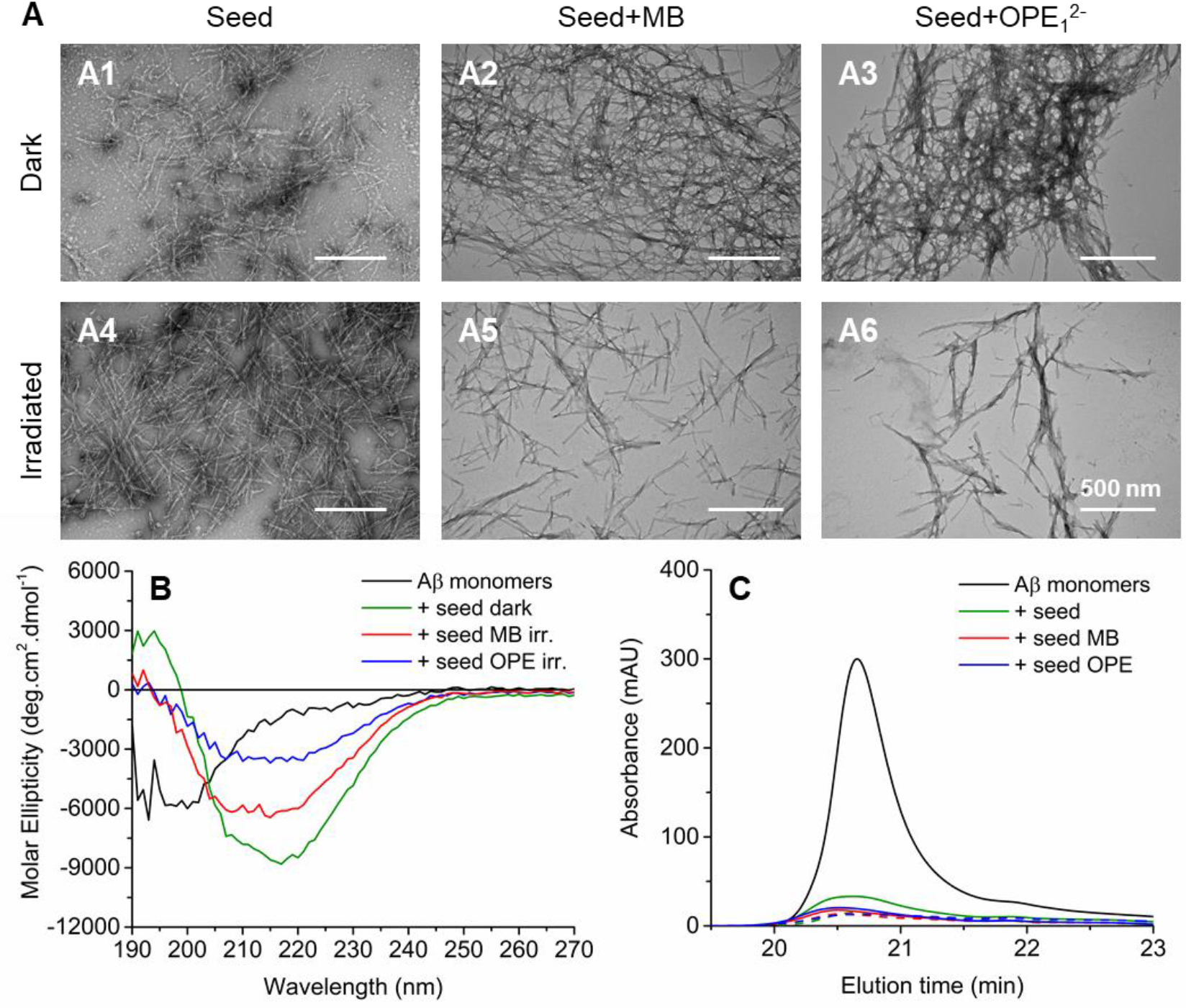
Oxidized fibrils largely retain the ability to seed Aβ monomer aggregation. **A**: TEM images of 50 µM Aβ40 monomers incubated with 2.6 µM non-oxidized (dark) or oxidized (irradiated) fibril seeds for 72 hours at 37 °C. Seeds were prepared by sonicating fibrils treated with MB or OPE_1_^2-^ (5 to 1 Aβ to photosensitizer molar ratio) in the dark or under irradiation for 4 hours. **B**: CD spectra of Aβ40 monomers before incubation (black) and after 72 hours of incubation in the presence of non-oxidized fibril seeds (green) or oxidized fibril seeds generated by MB (red) or OPE_1_^2-^ (blue). **C**. SE-HPLC chromatograms of Aβ40 monomers before incubation (black) and after 72 hours of incubation in the presence of non-irradiated fibril seeds (green) or fibril seeds in the presence of MB (red) or OPE (blue). Fibril seeds were either kept in the dark (solid line) or irradiated for 4 hours (dashed line) before seeding Aβ40 monomers. For HPLC, samples were first centrifuged to remove any insoluble aggregates before injecting onto the size exclusion column.

**Figure 10:**
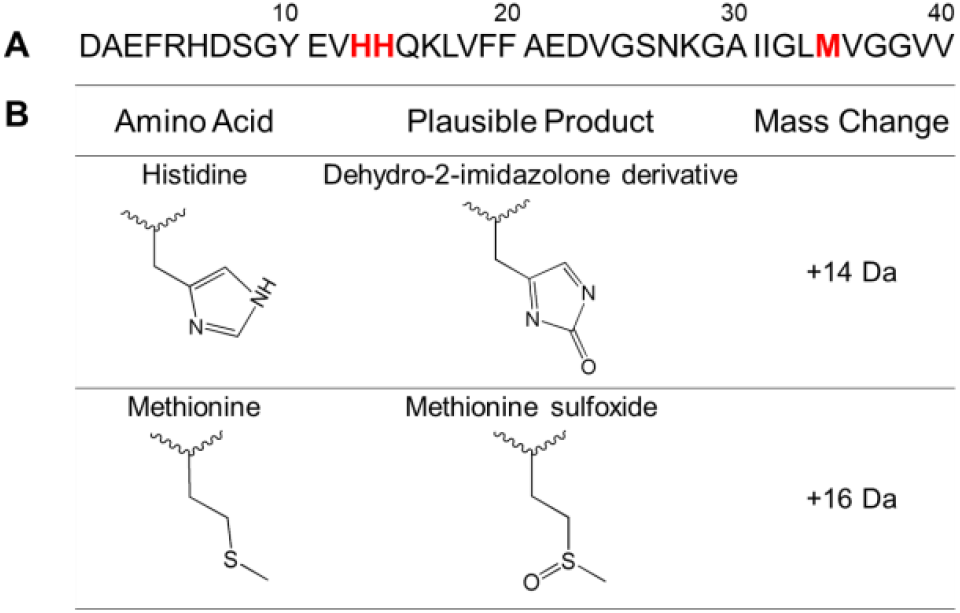
Photo-oxidation sites on Aβ40 (A) and plausible products of oxidation (B).

As shown in the TEM images, Aβ40 monomers formed large clusters of fibrils after 72 hours of incubation in the presence of fibril seeds or fibrils seeds that were irradiated for 4 hours (Figure 10A1 and 10A4). When the seeds were exposed to MB or OPE_1_^2-^ in the dark, the seeding potency was not affected as similar mature fibrils were observed after 72 hours of incubation (Figure 10A2 and 10A3). Fibril formation still occurred after incubating the monomers with oxidized seeds, either by MB or OPE. However, the fibrils produced are shorter and no large fibrillar clusters were observed (Figure 10A5 and 10A6). These fibrils were further characterized by CD spectroscopy. As shown in Figure 10B, fibrils produced in the presence of non-oxidized seeds (green) were rich in β-sheet as evidenced by the presence of a negative and positive CD signals at around 195 nm and 218 nm, respectively. Fibrils seeded by oxidized fibrils (MB in red and OPE_1_^2-^ in blue) also contained β-sheets. However, the CD signal was weaker. The percentages of Aβ40 monomers remaining after seeded incubation were determined by SEC (Figure 10C). About 18% monomers remained in the samples seeded with non-irradiated and irradiated seeds (Figure 10A1 and A4). Samples incubated with photo-oxidized seeds (Figure 10C: MB: dashed red line and OPE_1_^2-^: dashed blue line) contained about 10% monomers after 72 hours of incubation. Taken together, photo-oxidized fibrils did not show reduced seeding potency, but produced fibrils that were shorter and contained less β-sheets.

### Cell toxicity of oxidized Aβ40 fibrils

We have shown that both photo-oxidation by OPE_1_^2-^ and MB produced shorter fibrils that are still rich in β-sheets and able to seed further aggregation. It is essential to additionally investigate the effect of oxidation on fibril neurotoxicity as smaller oligomers and protofibrils are widely believed to be more toxic than mature fibrils. Furthermore, whether the photosensitizers are toxic to cells also need to be evaluated.

Toxicity of OPE_1_^2-^ and MB on SHSY-5Y neuroblastoma cells was evaluated at different concentrations (1-10 µM) both in the dark and with a 5-min irradiation. After treatment, cells were incubated at 37°C in 5% CO_2_ for 24 hours, after which cell viability was determined by the MTS assay. As shown in Figure 11A, viability of cells was not affected by the light treatment alone. When cells were treated with various concentrations of either MB or OPE_1_^2-^ and kept in the dark, cell viability was close to 100% indicating that both compounds were not cytotoxic at concentrations up to 10 µM. After exposure to light in the presence of varying concentrations of MB, cell viability was lower than the 70% threshold generally considered for cytotoxicity^72^ at MB concentrations higher than 5 µM. Note that cell viability was significantly lower in the irradiated samples compared to those kept in the dark (p-value ≤ 0.01). These results indicate that MB is not toxic for up to 10 µM, but becomes cytotoxic when exposed to light at concentrations higher than 5 µM. This cytotoxicity is likely due to light induced singlet oxygen generation by MB, which induced non-specific oxidation and led to cell death. In contrast, OPE_1_^2-^ did not reduce cell viability after light treatment even at 10 µM. The lack of toxicity indicates that OPE did not exhibit photosensitizing activity in the presence of the neuroblastoma cells. Given that OPE’s photosensitizing activity is turned-on by binding, this result also indicates a lack of non-specific binding of the OPE to the cells. Results from this assay point to the potential of the OPE as a selective, in addition to being controllable, photosensitizer with minimal off-target oxidation and toxicity.

**Figure 11:**
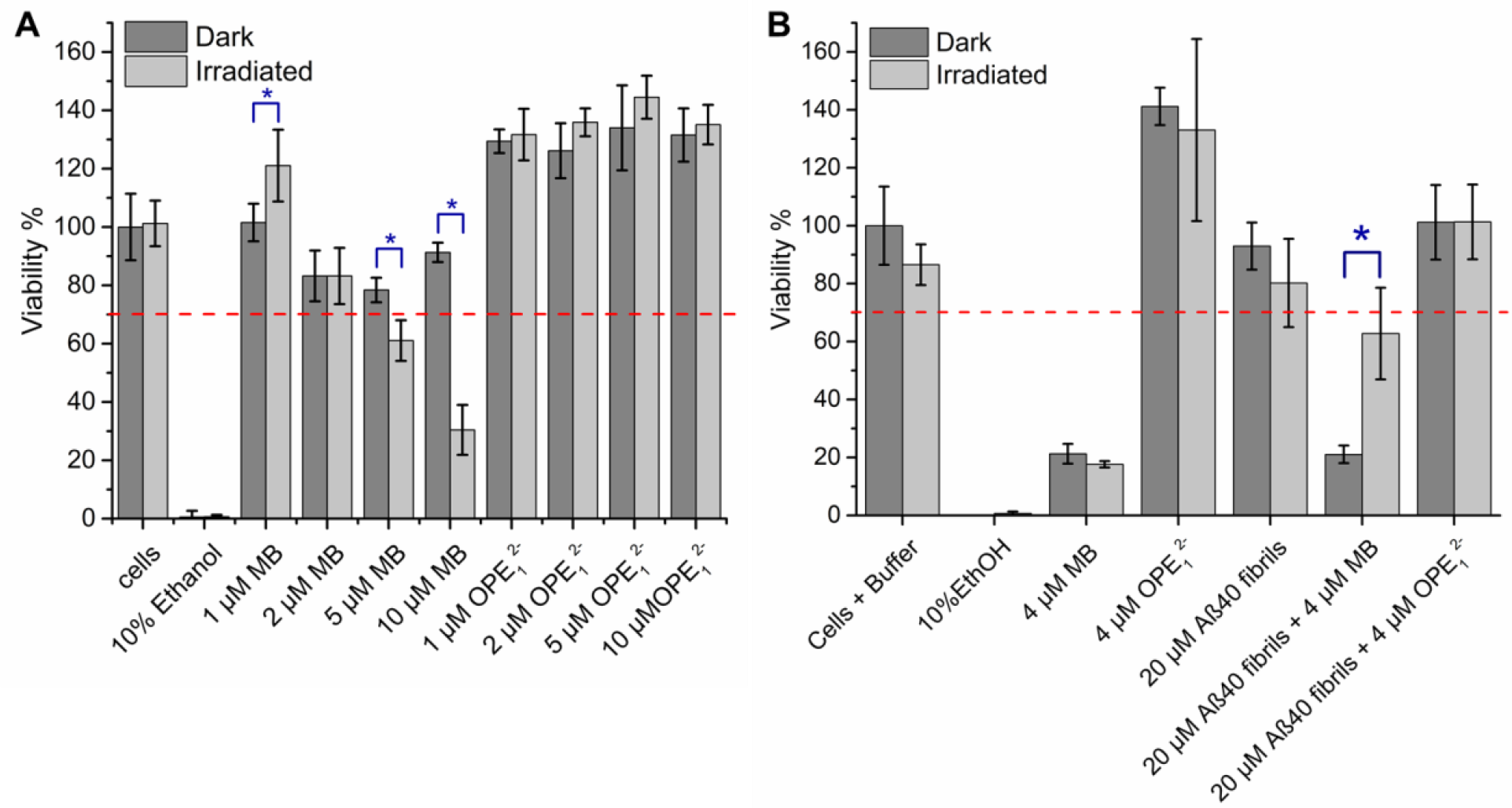
OPE_1_^2-^ and oxidized fibrils are not cytotoxic. (A) Viability of SHSY-5Y neuroblastoma cells incubated for 24 hr in the presence of varying concentrations of OPE_1_^2-^ or MB in the dark or after 5 min irradiation. (B) Viability of SHSY-5Y neuroblastoma cells incubated for 48 hr in the presence of MB, OPE_1_^2-^ or oxidized Aβ40 fibrils. In the irradiated condition, samples were exposed to light for 4 hrs prior to adding to SHSY-5Y cells. Cell viability was normalized to the negative control of untreated cells. Error bars represent standard deviations of quintuplet experiments. Red dashed line represents 70% viability threshold commonly used to define cytotoxicity^72^. Blue asterisks indicate significant differences between the dark and irradiated incubations (t-test with a p-value ≤ 0.01).

The effect of photo-oxidation on Aβ40 fibril toxicity was also investigated (Figure 11B). Fibrils were incubated at 50 µM in the dark or under irradiation for 4 hours in the absence or presence of 10 µM OPE_1_^2-^ or MB. After the 4-hour incubation, fibrils were added to SHSY-5Y cells at 20 µM with 4 µM photosensitizer. The cells were then incubated for 48 hours at 37 °C in 5% CO_2_ and cell viability was determined by MTS assay. As additional controls, cells were treated with 4 µM MB or OPE_1_^2-^ and incubated for 4 hours in the dark or under irradiation. In the presence of 4 µM MB, cell viability decreased by 80% for both dark and light conditions, indicating that MB is cytotoxic with a longer incubation period. Toxicity of MB might be caused by singlet oxygen generation when the samples were exposed to light during preparation. In contrast, the presence of 4 µM OPE_1_^2-^ did not cause any reduction in cell viability in both dark and light conditions, which further confirms that the OPE is not cytotoxic even after 48 hours of incubation. Irradiated and non-irradiated Aβ40 fibrils were also not cytotoxic. However, the standard deviation of cell viability in the presence of irradiated fibrils was larger (∼15% vs. ∼8%) which could be caused by the shorter fibrils produced with irradiation. Cells exposed to fibrils oxidized by OPE_1_^2-^ displayed similar viability compared to the non-irradiated fibrils indicating that the shorter oxidized fibrils as determined by TEM in Figure 9 were not cytotoxic.

Interestingly, cells treated with fibrils incubated with MB in the dark showed around 20% viability, comparable with cells treated with just MB. These results are consistent in that fibrils are not toxic and the toxicity in these samples are likely caused by short light exposure during sample preparation that activated MB photosensitizing activity. Surprisingly, cells treated with fibrils irradiated with MB showed a higher ∼60% viability. Although the exact cause for this is not known, we hypothesize that the lower toxicity of the irradiated samples is due to MB photo-bleaching or photodegradation during the 4 hours of irradiation, discoloration of the sample was observed after irradiation. Note that in both sets of SHSY-5Y viability experiments, greater than 100% viability was consistently observed when the cells were treated with OPE, although the cause of this increased viability is not yet clear.

## Discussion

PDT is an attractive method to photo-oxidize amyloid aggregates, which could promote their disassembly and clearance to treat neurodegenerative disorders such as Alzheimer’s disease. The feasibility of such an approach hinges on the discovery of photosensitizers that are selective for the pathogenic, aggregated conformations of amyloid proteins to avoid off-target oxidation that can lead to the loss-of-function of the native amyloid proteins or the death of surrounding cells or tissue. Additionally, the photosensitizer needs to be non-toxic, as do their photo-oxidized products. In this study, we demonstrate that a novel phenylene ethynylene-based oligomer OPE_1_^2-^ selectively and controllably photo-oxidizes the fibrils, but not the monomers, of Aβ40. The oxidized fibrils retain its β-sheet rich structures and fibril-seeding ability, and are nontoxic.

### OPE_1_^2-^ selectively and controllably photo-oxidizes Aβ40 fibrils over monomers

Three techniques were used to thoroughly evaluate the oxidation of Aβ40 monomers and fibrils exposed to light in the presence of OPE_1_^2-^ or the well-known, but nonselective MB. DNPH dot-blot (Figure 3), ESI-MS (Figures 4 and 5), and amino acid analysis (Figure 6) results revealed that OPE_1_^2-^ is a light-controllable photosensitizer that selectively oxidizes Aβ40 fibrils over its monomeric counterpart with minimal off-target oxidation. The selective oxidation of the fibrillar conformation can be explained by the high binding affinity of OPE_1_^2-^ to the fibrils (K_d_ = 0.70 ± μM)^52^ and its weak interaction with the monomers as shown by a lack of OPE fluorescence turn-on (Figure 2B). In contrast, MB oxidizes both Aβ40 monomers and fibrils under irradiation. This lack of selectivity of MB may be attributed to its non-specific interactions to the negatively charged Aβ in both conformations as MB has been previously reported to interaction with monomeric Aβ42 with a dissociation constant K_d_ of 48.7 ± 3.6 μM.^44^

### OPE_1_^2-^ photo-oxidizes His13, H14 and Met35 residues in Aβ40 fibrils

In Aβ40, five amino acids can be photo-oxidized: 3 histidines (His6, His13, His14), 1 tyrosine (Tyr10) and 1 methionine (Met35)^59^. Amino acid analysis (Figure 6) and mass spectrometry (Figure 5) showed that light treatment in the presence of OPE_1_^2-^ or MB led to the oxidation of 2 His and 1 Met in Aβ40 fibrils. Interestingly, OPE_1_^2-^ only partially oxidized Met35 which might be due to a lower singlet oxygen quantum yield compared to that of MB, which is about 0.5.^46^ However, as Met35 is located in the core of the fibril (Figure 8), its accessibility to singlet oxygen and subsequent reaction oxygen species may be reduced compared to fibril-surface exposed His13 and His14. To better understand the difference between the photosensitizing activities of OPE_1_^2-^ and MB, their singlet oxygen quantum yields could be characterized by luminescence or photochemical methods^73^. Interestingly, Tyr10 in fibrils is not oxidized by either photosensitizer despite its close proximity to His13 and His14 which were both oxidized. The low propensity of tyrosine to be photo-oxidized may be due to its lower rate constant (K) for ^1^O_2_ quenching compared to histidine and methionine (K_His_ = 4.6 × 10^−7^ > K_Met_ = 1.3 × 10^−7^ > K_Tyr_ = 0.2-0.5 × 10^−7^ M^-1^s^-1^)^67,74^. Moreover, photo-oxidation of His13, His14, and Met35 residues in Aβ40 fibrils observed in this study (Figure 12) has also been previously reported in a study where a flavin-based catalyst and an iridium (III) complex were used^35,75^.

### OPE_1_^2-^ photo-sensitization disassembles Aβ fibrils into β-sheet rich, non-toxic, and seeding-competent protofibrils

Aβ40 fibrils are highly thermodynamically stable^76^, which makes their degradation challenging. Photo-oxidation of Aβ40 fibrils reduces their overall hydrophobicity and could promote their breakdown^77^. To better understand the effect of oxidation on fibril morphology and secondary structures, we analyzed oxidized fibrils by TEM imaging and CD spectroscopy (Figure 9), respectively. Results show that irradiation by either MB or OPE_1_^2-^ broke clusters of long Aβ40 fibrils into shorter fibrils still rich in β-sheets. Fibril disassembly could be caused by the partial oxidation of Met35 located in the core of the fibril (Figure 7C), which stabilizes the long fibrils^77^. As protofibrils and oligomers are known to be more toxic than fibrils^9-11,78^, we evaluated cell toxicity of the oxidized fibrils (Figure). We observed that oxidized fibrils do not display higher toxicity compared to non-oxidized fibrils indicating that using PDT to break down the fibrils will not generate more toxic species. Additionally, OPE_1_ ^2-^ is not cytotoxic to cells, even under irradiation, which further supports their use in PDT. The seeding potency of oxidized fibrils was also characterized (Figure C). Oxidized fibril seeds maintained their seeding capacity, however they produced shorter fibrils that were less rich in β-sheet compared to mature non-oxidized fibrils.

## Conclusions

Oligomeric conjugated polyelectrolytes such as OPE_1_^2-^ have been recently shown to selectively and sensitively bind to aggregated conformations of a number of amyloid proteins over their monomeric conformers.^52^ Concomitant with aggregate binding is the turn-on of OPE’s fluorescence and singlet oxygen generation,^57^ suggesting that OPEs are potentially superior photosensitizers for the PDT treatment of protein misfolding diseases such as Alzheimer’s disease. In this study, we carry out a proof-of-concept investigation of the controlled and selective photo-oxidation of Aβ40 fibrils over monomers by OPE_1_^2-^ and compared its photosensitizing activity to MB. We showed that MB non-selectively oxidizes both Aβ monomers and fibrils, while OPE_1_^2-^ only oxidizes Aβ fibrils. Three amino acids on the fibril are oxidized by OPE photosensitization, His13, His14 and Met35, which proceeds through binding induced generation of ^1^O_2_. Photo-oxidation causes fibrils to disassemble into shorter, but non-toxic oxidized fibrils. The oxidized fibrils also retain their ability to seed further Aβ40 aggregation, albeit fibrils of a lower β-sheet content were produced. Overall, this study demonstrates the ability of OPE_1_^2-^ to controllably and selectively photo-sensitize the oxidation of Aβ fibrils. The selective nature of OPE’s photosensitizing activity overcomes the major drawback of off-target oxidation from using conventional photosensitizers such as MB in PDT. Combined with its selective fluorescent sensing capabilities, our results from this study support the further development of OPEs as potential theranostics for the simultaneous detection and clearance of amyloid aggregates in protein misfolding diseases such as Alzheimer’s disease.

## Experimental Methods

### Materials

Synthetic amyloid-β (1-40) (Aβ40) was purchased from Peptide 2.0 (Chantilly, VA). Tris was obtained from BioRad (Hercules, CA). Sodium chloride (NaCl), dimethyl sulfoxide (DMSO), sodium azide, acetonitrile, methanol and hydrochloride acid (HCl) were acquired from EMD Millipore (Burlington, MA). SH-SY5Y neuroblastoma cells, Dulbecco’s Modified Eagle’s Medium (DMEM) F12 media, fetal bovine serum (FBS), 2,4-dinitrophenylhydrazine (DNPH), trifluoroacetic acid (TFA), and Tween® 20 were purchased from Sigma-Aldrich (St. Louis, MO). Penicillin-streptomycin (PS) at 10,000 U/mL, AP Rabbit anti-Goat IgG (H+L) secondary antibody and 1-Step NBT-BCIP substrate were purchased from Thermo Fisher (Waltham, MA). The CellTiter 96® AQueous One Solution Cell Proliferation Assay was purchased form Promega (Madison, WI). Goat anti-DNP primary antibody was acquired from Bethyl (Montgomery, TX). OPE_1_^2-^ was synthesized and purified by previously published procedures^55^. MB was purchased from Avantor (Radnor, PA). 400 mesh copper grids covered by a Formvar/Carbon film (5-10 nm) were obtained from Ted Pella (Redding, CA) and 2% aqueous uranyl acetate was purchased from Electron Microscopy Sciences (Hatfield, PA).

### Aβ40 monomers and fibrils preparation

Lyophilized Aβ40 peptide was solubilized in DMSO at 50 mg/mL. After centrifugation at 14,000 rpm for 15 min, the supernatant was removed and stored at -70 °C. Monomeric Aβ was prepared by diluting the stock solution to 150 µM with a pH 8.0 40 mM Tris buffer containing 150 mM NaCl and 0.01% sodium azide. Fibrils were made by incubating Aβ monomers at 37 °C for 23 days^52^. Photo-oxidation of both Aβ40 monomers and fibrils by either OPE_1_^2-^ or MB was performed by incubating the peptides with a photosensitizer at a peptide to photosensitizer molar ratio of 5 to 1 or 25 to 1. The mixtures were either kept in the dark or exposed to light in a photochamber (Luzchem Research Inc.) using 10 LZC 420 lamps (Osram Sylvania, Wilmington, MA) which emitted UV light between 350 and 700 nm at 8 W per lamp. Samples were then irradiated for 0-6 hours for the various experiments described below.

### Seeding experiment preparation

Stock Aβ at 50 mg/mL in DMSO was diluted to 50 µM using 50 mM phosphate buffer (PB) with 100 mM NaCl at pH 7.4. This Aβ40 monomer solution was either incubated alone or in the presence of 2.6 µM Aβ40 seed protofibrils at 37 °C for 72 hours. Three different seed protofibrils were prepared from mature fibrils: (1) Aβ40 fibrils incubated for 4 hours in the dark or under irradiation, (2) Aβ40 fibrils incubated in the presence of OPE_1_^2-^ for 4 hours (5 to 1 molar ratio Aβ to photosensitizer) in the dark or under irradiation and (3) Aβ40 fibrils incubated in the presence of MB for 4 hours (5 to 1 molar ratio Aβ to photosensitizer) in the dark or under irradiation. Irradiation was carried out in a photo-chamber (Luzchem Research, Inc.) containing 10 bulbs (350-750 nm). Seed protofibrils were then prepared by sonicating the above incubated fibrils for 10 minutes using a 550T Ultrasonic Cleaner (VWR International, Radnor, PA). Note that the mature fibrils prepared by 23-day incubation were not processed, i.e., washed or undergone any separation, prior to the 4 hour incubation and sonication to produce seed protofibrils.

### Absorbance and fluorescence measurements

OPE_1_^2-^ and MB absorbance and emission spectra were recorded at 1 µM in the presence of varying concentrations of Aβ40 (0, 1, 3 and 5 µM) in pH 7.4 10 mM PB after 30 minutes of incubation in the dark at room temperature. Absorbance spectra were obtained with a Lambda 35 UV/VIS spectrometer (PerkinElmer, Waltham, MA) in a quartz cuvette (PerkinElmer, Waltham, MA). Emission scans were obtained at excitation wavelengths of 390 nm and 660 nm for OPE_1_^2-^ and MB, respectively, and were recorded using a PTI QuantaMaster 40 steady state spectrofluorometer (HORIBA Scientific, Edison, NJ) in a quartz cuvette (Starna cells Inc., Atascadero, CA).

### DNPH dot blot

µm PVDF membrane (ThermoFisher, Waltham, MA) was soaked in 100% methanol for 15 seconds, then in water for 5 minutes, and finally in pH 7.4 phosphate buffered saline containing 0.1% Tween 20 (TPBS) for 15 min. After incubating Aβ40 monomers or fibrils with a photosensitizer (5 µM protein with 1 µM photosensitizer) in the dark or under illumination for up to 6 hours, samples were blotted onto the membrane four times at 1 µL each and let dry for 15 minutes. DNPH derivatization of protein carbonyl groups was carried out as previously described^79^. Briefly, the membrane was equilibrated in 2.5 N HCl for 5 minutes before transferring to a 20 mM DNPH solution in 2.5 N HCl for 5 minutes. Excess DNPH was then washed away with three 5 mL aliquots of 2.5 N HCl and then 5 mL of 100% methanol. The membrane was then immuno-stained. First the membrane was submerged in the blocking buffer (pH 7.4 phosphate buffered saline (PBS) containing 5% nonfat dry milk and 0.1% Tween 20) for 24 hours at room temperature. The membrane was then washed six times with a washing buffer (PBS containing 0.1% Tween 20) for 5 min per wash before applying the goat anti-DNP primary antibody diluted at 1:10,000 in blocking buffer for 2 hours under agitation in the dark. The membrane was then washed 6 times with the washing buffer before applying the rabbit anti-goat IgG secondary antibody, alkaline phosphatase (AP) conjugate diluted at 1:10,000 in blocking buffer, for 2 hours under agitation in the dark. The membrane was washed 3 time with the washing buffer and 3 time with PBS before revealing the dot blot with the 1-Step NBT-BCIP substrate. Once the dots appeared, the membrane was rinsed twice with distilled water for 2 minutes each under agitation and was dried overnight before imaging the membrane.

### Amino acid analysis (AAA)

Samples containing 25 µM Aβ40 and 5 µM photosensitizer were kept in the dark or were irradiated for 4 hours. After light treatment, samples were sent to the Molecular Structural Facility at University of California Davis for amino acid analysis (AAA) using a sodium citrate buffer system. Briefly, this analysis consisted of drying 100 µL of 25 µM peptide samples and performing a liquid phase hydrolysis using 200 µL 6 N HCl containing 1% phenol for 24 hours at 110 °C. After hydrolysis, the protein was dried and added to norleucine, an internal standard, to reach a final volume of 200 µL. The samples were analyzed on a cation-exchange chromatography column using a L-8800 Hitachi analyzer and a post column ninhydrin reaction detection system.

### Electrospray ionization mass spectrometry (ESI-MS)

Both monomeric and fibrillar proteins were desalted first using the Amicon Centrifugal Filter (Millipore Sigma, Burlington, MA) before analyzed by ESI-MS. 250 µL of peptide at 150 µM was loaded onto the filter to which 4 mL of PB was added before centrifuging the filter at 3500 rpm for 20 min. After four washing steps using PB buffer, the retentate was collected and volume adjusted to 250 µL. The protein concentration was then determined using the Bradford protein concentration assay (Sigma-Aldrich, St. Louis, MO).

Desalted Aβ40 monomer and fibril samples were analyzed by ESI-MS after 4 hours of irradiation in the absence and presence of MB or OPE_1_^2-^ (25 µM protein with 1 µM photosensitizer). Before analysis, Aβ40 fibrils were digested using the Endoproteinase Lys C enzyme (New England BioLabs, Ipswich, MA) at a 1/50 (w/w) enzyme to protein ratio. The digestion was performed by incubating the samples at 37 °C for 16 hours. Digested Aβ40 monomers and fibrils were diluted to 5 µg/mL using acetonitrile with 1% TFA and were analyzed under continuous ESI-MS spray on SYNAPT G2 Mass Spectrometer. This analysis was performed in a positive mode by using the following settings: Capillary = 3.5 kV, sampling cone = 251, extraction cone = 5, source temperature = 120 °C, desolvation temperature = 300 °C, and desolvation gas flow = 650 L/h. Data were analyzed with the software MassLynxV4.1 for generation of calculated and comparison to observed mass ion packet.

### Size exclusion chromatograph (SEC)

The unincubated and incubated Aβ40 samples were analyzed by SEC to quantify the amounts of monomers present in the samples after 72 hours of incubation. Before injecting the sample on the HPLC column, 60 µL of 50 µM Aβ40 was centrifuged for 15 min at 14,000 rpm to remove insoluble aggregates. Supernatant (50 µL) was injected onto a BioSec-SEC-s2000 (Phenomenex, Torrance, CA) column that was already equilibrated with 10 mM phosphate buffer saline (PBS) at pH 7.4 at 0.5 mL/min on an Agilent 1100 series HPLC (Agilent Technology, Santa Clara, CA). Absorbance at 215 nm was monitored. Background signal was subtracted using the Agilent ChemStation software and percentages of soluble monomers were calculated relative to the protein present in the unincubated samples.

### Circular dichroism (CD) spectroscopy

Desalted protein solutions were diluted to 50 µM in 10 mM PB in the absence or presence of 10 µM MB or OPE_1_^2-^. After irradiation, the peptide solution was loaded into a quartz cuvette with a path length of 1 mm (Starna cells Inc, Atascadero, CA) and analyzed on an AVIV 410 CD Spectrometer (AVIV, Lakewood, NJ) between 190 and 270 nm using an averaging time of 15 seconds. Three scans were recorded per sample and averaged signal was converted to molar ellipticity^80^.

### TEM imaging

Aβ40 samples were diluted to 5 µM using MilliQ water. After the grids were glow discharged (Harrick Plasma Cleaner, Carson City, NV) for 30 seconds, each sample was loaded onto a grid and let adsorbed for 5 min. After wicking away excess sample, the grid was stained one time for 3 minutes and three times for 1-minute each using 2% uranyl acetate (Electron Microscopy Sciences, Hatfield, PA). Excess stain was wicked away in between the steps. The grid was then air dried for 30 minutes and imaged using a HITACHI HT7700 transmission electron microscope (Hitachi High Technologies Corp., Tokyo, Japan) with a beam current of 8.0 µA and an accelerating voltage of 80 keV.

### Cell toxicity assay

Neuroblastoma SHSY-5Y cells were cultivated in DMEM media containing 1% PS and 10% FBS at 37°C and 5% CO_2_. When the cells reached 80% confluency, they were used to set up 96 well plates with 20,000 cells/100 µL well. After 16 hours of incubation, media was changed with serum deprived DMEM media and cells were further incubated for 24 hours to ensure cell synchronization. Cells were then treated with a sample containing appropriate protein and sensitizer concentrations and were incubated for another 24 or 48 hours after which cell viability was monitored by MTS assay. The assay involved adding 20 µL MTS reagent to each well already containing 100 µL media and incubating the plate for 3 hours before measuring the absorbance at 490 nm with a Spectra Max M2 plate reader (Molecular Devices, Sunnyvale, CA). The background absorbance of MTS in media alone was subtracted from the absorbance obtained in the presence of cells. Absorbance obtained for untreated cells were also obtained and used as 100% viability.

### All atom molecular dynamics (MD) simulation

The initial configurations for the Aβ fibril - OPE_1_^2-^ simulations were built with UCSF Chimera^81^. The fibril structure, 2LMN^68^ was obtained from the Protein Data Bank. In this work, we prepared an Aβ9-40 protofibril made of 24 peptides to evaluate OPE_1_^2-^ binding sites. 12 OPEs were positioned around the protofibril at a distance of 10 Å away from the protofibril surface. The anionic OPE_1_^2-^ was built using the GaussView 5 package and geometry optimizations was carried out with Gaussian 09.^82^ Simulations were prepared using the AMBERTools suite^83^. Parameters for simulating the protofibril structures were obtained from the AMBER14 force field. OPEs were parameterized using the AMBER generalized force field (GAFF) and partial charges were generated using the R.E.D. server^84,85^. Each system was solvated in explicit water molecules using the TIP3P model and a total of 48 counter Na^+^ ions were added to neutralize the system. MD was performed using the AMBER MD package as previously described^86^. Coordinates for the single Aβ9-40 monomer hairpin were extracted from the protofibril PDB structure following energy minimization and equilibration of the monomer peptide. Production MD simulations were carried out for 100 ns.

OPE_1_^2-^ binding sites on the protofibril surface were analyzed using UCSF Chimera to determine the proximity of bound OPEs to oxidizable residues on the protofibrils. Residues within 4 Å of bound OPEs (either single or complexed) were selected and counted. Distances of the oxidizable methionine residues to bound OPEs, which were larger than the 4 Å cutoff, were determined as the center-of-mass distances between methionine residues and bound OPEs.

## ASSOCIATED CONTENT

### Supporting Information

Reverse phase HPLC of Aβ40 monomers before and after irradiation in the presence of MB and OPE _1_^2-^; TEM images of Aβ40 after incubation of 50 µM monomers either alone (A) or in the presence of 2.6 µM non-sonicated Aβ40 fibrils (B) or 2.6 µM sonicated Aβ40 fibrils (C).

## AUTHOR INFORMATION

### Author Contributions

A.M.F., D.O., F.A.M, T.D.M., J.S.M, D.G.E., D.G.W., and E.Y.C designed research; A.M.F.,D.O., F.A.M., J.H., F.M, T.D.M., G.B., and L.M.P. performed research; A.M.F., D.O., F.M, J.S.M., D.G.E. and E.Y.C. analyzed data; A.M.F. and E.Y.C. wrote the paper.

## ACKNOWLEDGMENT

This work was performed, in part, at the Center for Integrated Nanotechnologies, an Office of Science User Facility operated for the U.S. Department of Energy (DOE) Office of Science. Los Alamos National Laboratory, an affirmative action equal opportunity employer, is managed by Triad National Security, LLC for the U.S. Department of Energy’s NNSA, under contract 89233218CNA000001. We are also thankful to Dr. Yanli Tang and Dr. Eunkyung Ji who originally synthesized OPE_1_^2-^. The Extreme Science and Engineering Discovery Environment (XSEDE), which is supported by National Science Foundation grant ACI-1053575, was used for performing simulations.

## ABBREVIATIONS

AAA: amino acid analysis
Aβ: amyloid β
CD: circular dichroism
CTAB: cetyltrimethylammonium bromide
DMEM: Dulbecco’s Modified Eagle’s Medium
DMSO: dimethyl sulfoxide
DNPH: 2,4-Dinitrophenylhydrazine
ESI-MS: electrospray ionization mass spectrometry
FBS: fetal bovine serum
GAFF: generalized force field
HCl: hydrochloride acid
RP-HPLC: reverse phase high performance liquid chromatography
MB: methylene blue
MD: molecular dynamics
NaCl: sodium chloride
OPE: oligo-p-phenylene ethynylene
PDB: protein databank
PDT: photodynamic therapy
PS: penicillin-streptomycin
TPBS: pH 7.4 phosphate buffered saline containing 0.1% Tween 20
TEM: transmission electron microscopy
TFA: trifluoroacetic acid

## Table of Content

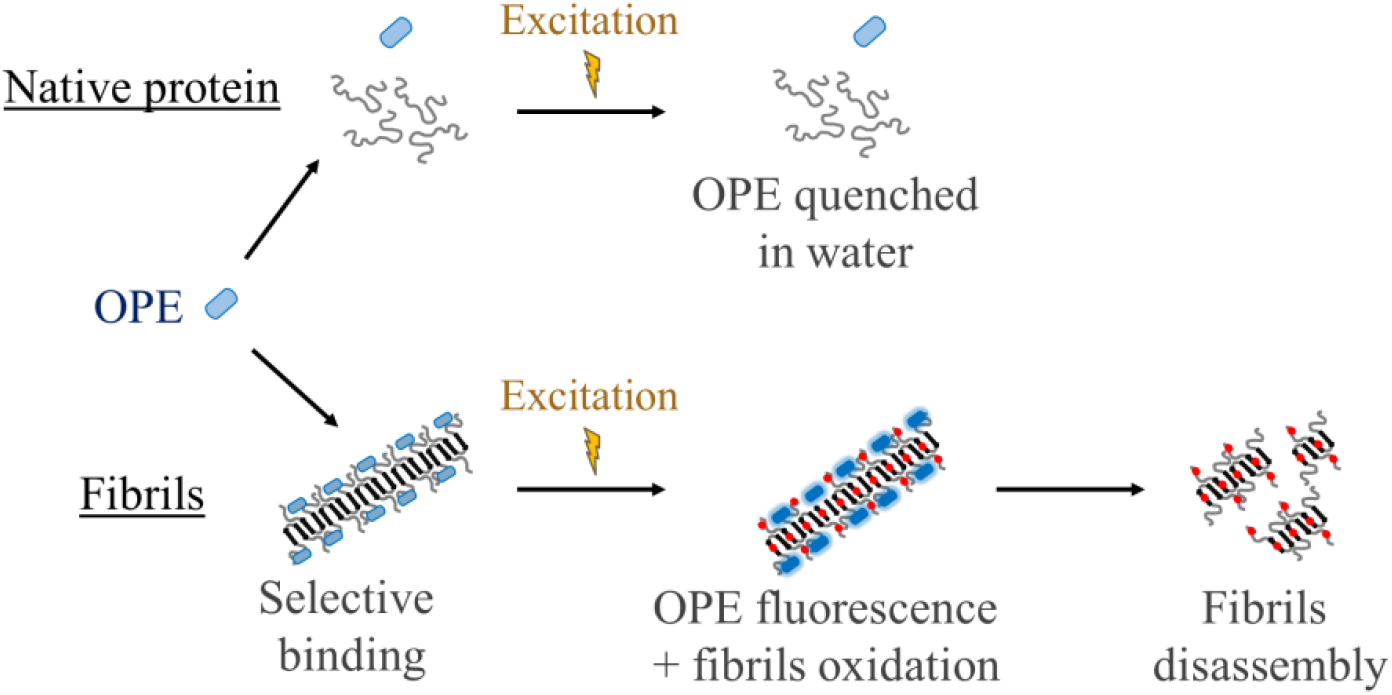

## Supplementary Information

**Figure S1:**
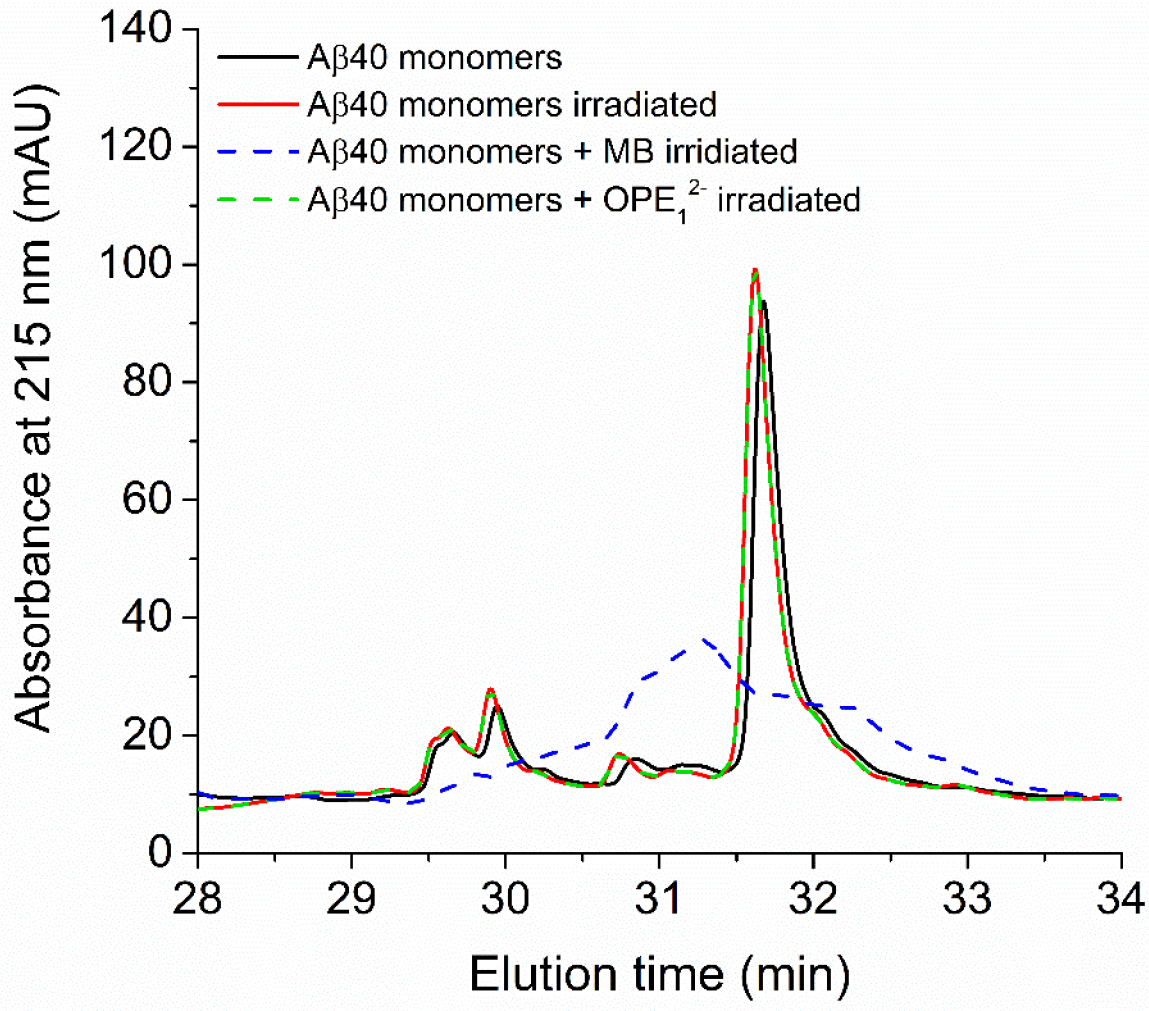
Reverse phase HPLC chromatographs of soluble Aβ40 before and after irradiation in the presence of MB or OPE_1_^2-^. Unincubated Aβ40 monomers, whether non-irradiated or irradiated in the presence of OPE_1_^2-^, display a similar elution time that corresponds to 44.6-44.4% acetonitrile. When Aβ40 monomers are irradiated in the presence of MB, the elution profile of Aβ40 monomers significantly changed. The main peak eluted earlier (43.5% acetonitrile), which shows that Aβ40 peptide became more hydrophilic, which is consistently with the oxidation of the peptide. Also, the elution profile is broad with multiple peaks, indicating the presence of a several populations of Aβ40. Experimental method: The monomeric protein was analyzed by RP-HPLC on an Agilent 1100 instrument (Agilent Technology, Santa Clara, CA) before and after irradiation in the presence of OPE_1_^2-^ or MB (5 µM protein with 1 µM photosensitizer). 110 µL of 5 µM protein was centrifuged at 14,000 rpm for 15 minutes. The supernatant (100 µL) was loaded onto an Eclipse XDB C18 column (Agilent Technology, Santa Clara, CA) pre-equilibrated at 40 °C with 95% of mobile phase A (water containing 0.1% TFA) and 5% of mobile phase B (acetonitrile containing 0.1% TFA). Aβ40 was eluted using a 5-100% linear gradient of mobile phase B over 40 min. The absorbance at 215 nm was monitored. Each chromatogram was background subtracted using the Agilent ChemStation software.

**Figure S2:**
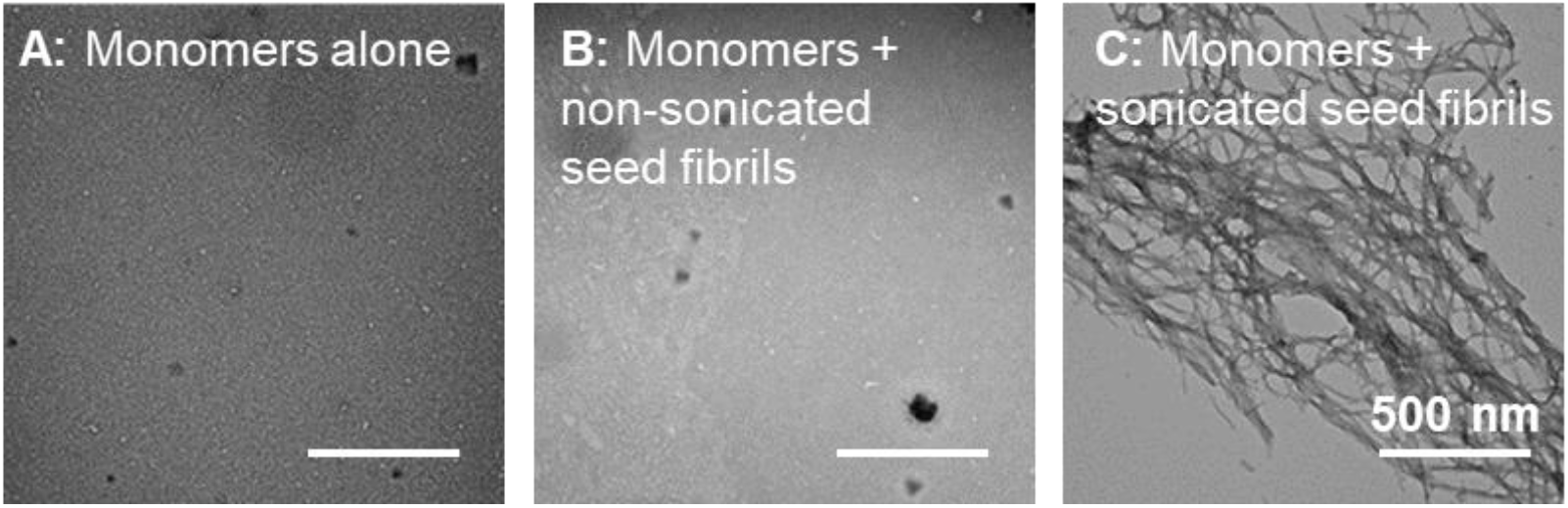
TEM images of Aβ40 after incubation of 50 µM monomers either alone (A) or in the presence of 2.6 µM non-sonicated Aβ40 fibrils (B) or 2.6 µM sonicated Aβ40 fibrils (C). Only Aβ protofibrils produced by sonication promoted fast peptide fibrillation.

